# Inferability of transcriptional networks from large scale gene deletion studies

**DOI:** 10.1101/082925

**Authors:** Christopher Frederik Blum, Nadia Heramvand, Armin S. Khonsari, Markus Kollmann

## Abstract

Generating a comprehensive map of molecular interactions in living cells is difficult and great efforts are undertaken to infer molecular interactions from large scale perturbation experiments. Here, we develop the analytical and numerical tools to quantify the fundamental limits for inferring transcriptional networks from gene knockout screens and introduce a network inference method that is unbiased and scalable to large network sizes. We show that it is possible to infer gene regulatory interactions with high statistical significance, even if prior knowledge about potential regulators is absent.

---

The functionality of a living cell is determined by the interplay of multiple molecular components that interact with each other. Generating a global map of these molecular interactions is an essential step to advance our understanding of the molecular mechanisms behind disease, development, and the reprogramming of organisms for biotechnological applications ^1^. The current advances in gene editing methods ^2^ have scaled up the size of genome-wide single and double knockout libraries, ranging from microbes ^3^,^4^ to higher eukaryotes ^5^ and open up a much more informative data source than inferring gene regulatory networks from unspecific perturbations, such as stress or changes in growth conditions ^6^. However, the detection of direct interactions between two genes from association measures – for example the covariance between transcript levels – remains a highly non-trivial task, given the significant noise among biological replicates, the frequent case where the number of parameters exceeds the number of independent data points, and the high dimensionality of the inference problem.

The existence of a direct interaction between gene A as a source of regulation (source node) and gene B as a target of regulation (target node) can be detected if a significant part of the transcriptional activity of B can be explained by the transcriptional activity of A but not by the activities of the remaining genes in the network. Thus a necessary condition for identifiability or inferability of links is the knowledge about the information that can be transmitted by alternative routes in the network, which can be obtained by perturbing the involved nodes ^7^. As most gene perturbation screens are incomplete – for example due to the fact that essential genes cannot be knocked out – we have in general the situation that a significant amount of interactions within an N-gene network are non-inferable, regardless of the amount of experimental replicates and the strength of perturbations. Direct interactions inferred from transcriptome data typically oversimplify the molecular complexity behind gene regulation, which frequently involves protein-protein interactions and modifications on protein or DNA level.

To calculate the maximum number of links that can be inferred from knockout screens in absence of other constraints, we consider a directed network of *N* nodes, with node activities as observables and a fraction *q* of node activities strongly perturbed by external forces. Given that the perturbed nodes are randomly distributed within the network, we can analytically calculate the expected fraction of inferable links, *F*(*q*), by a counting procedure illustrated in Fig. 1a (Online Methods and Supplementary Note 1).

**Figure 1.**
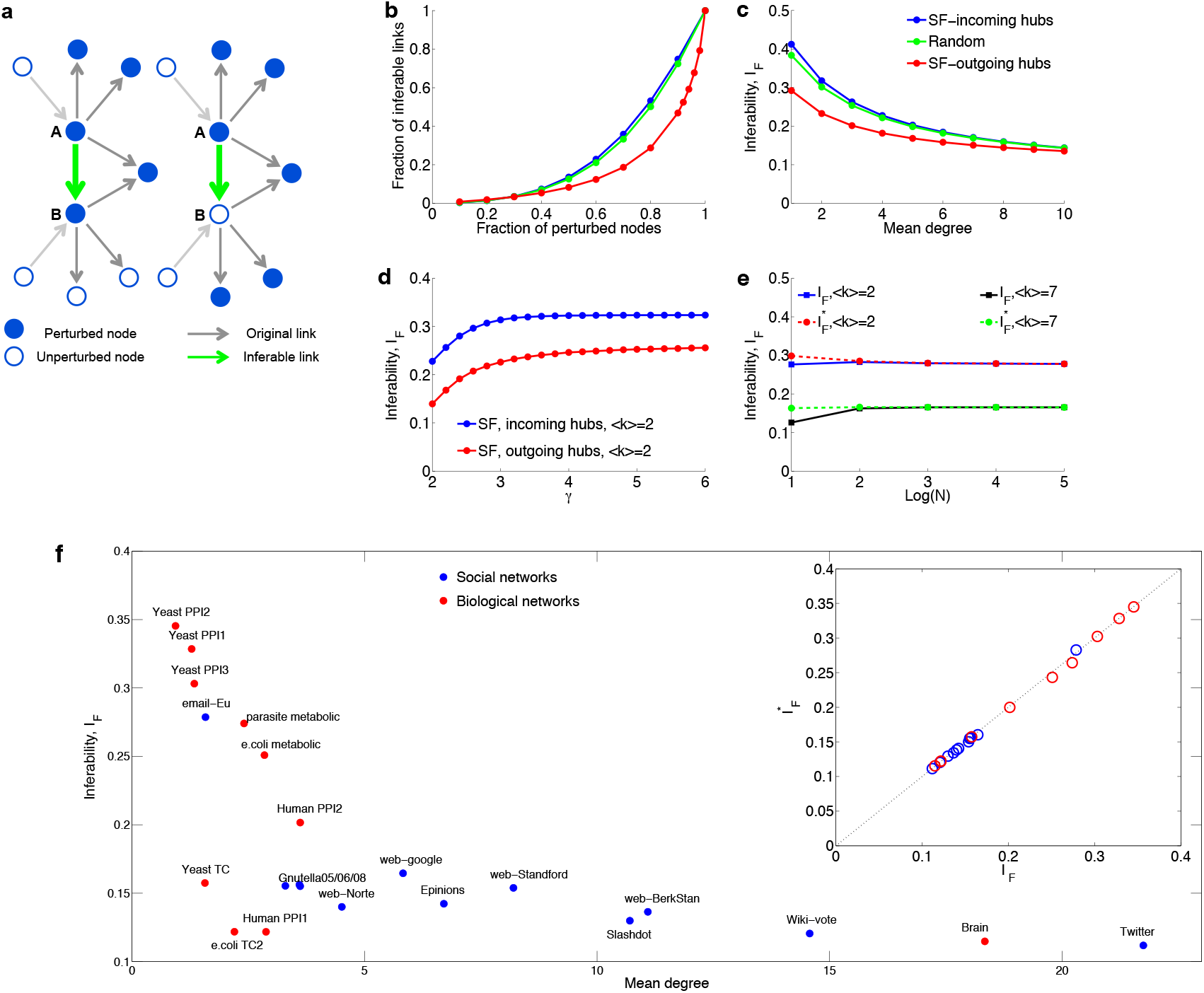
Inferability as a function of network parameters. **(a)** Directed links are inferable if either all outgoing links of the source node are perturbed including the target node (left panel) or if all outgoing links of source node and target node are perturbed, with the target node not perturbed. **(b)** Fraction of inferable links against the fraction of perturbed nodes using three network types: (i) scale free network with exponent *γ* = 2.5 and mean degree ⟨*k*⟩ = 3, where nodes of higher degree target nodes of lower degree (outgoing hubs), (ii) same network as (i) but with all link directions inverted (incoming hubs), or a network generated by random insertion of links with the same mean degree as scale-free networks (random network). Colour coding as in (c). **(c)** Network inferability versus mean degree, using networks of (b). **(d)** Network inferability versus scaling exponent for two types of scale-free networks. **(e)** Asymptotic invariance of the two inferability measures introduced in the main text with respect to network size. **(d)** Network inferability as a function of mean degree for social and biological networks. Correlation between the two inferability measures introduced in the main text (inset).

As *F*(*q*) is an upper bound for the expected number of directed links that can be inferred from stationary node activities, we now ask how this bound is related to the structural properties of the network. To compare different network architectures, it is useful to define the network inferability, *I*_*F*_, as the area under the *F*(*q*)-curve, 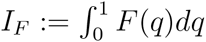. Comparison of *I*_*F*_ between two general classes of network structures with node degrees either power law or Poisson distributed shows that networks that are enriched with nodes of high out degree (outgoing hubs) are the most difficult ones to infer (Fig. 1b). Differences in inferability due to network structure are most predominant for networks with low mean degree and become less predominant with high mean degree (Fig. 1c). As our measure of inferablity, *I*_*F*_, is essentially determined by the outdegree distribution the curve starts saturating for scale-free exponents *γ* > 3, as in this regime the variance of the number of links per node is essentially constant for increasing *γ* and fixed mean degree ^8^ (Fig. 1d). The network inferability, *I*_*F*_, is asymptotically independent of network size (Fig. 1e). We further investigated the inferability of causal interactions in biological and social networks as a function of the mean degree (Fig. 1f). The decreasing trend can be explained by the higher number of alternative routes that come with a stronger connected network. The low inferability of gene regulatory networks can be attributed to master regulators that regulate a large fraction of the genome (hubs with high outdegree), whereas the comparatively high inferability of protein interaction networks is a consequence of the low number of different binding domains per protein and that only a fraction of the existing interactions have been identified due to limitations of experimental methods ^9^. If we assume that the probability *P*(*k, l,m\k → l*) can be factorized, the resulting inferability measure, 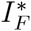, is simply a function of the outdegree distributions, *P*(*k*) and *P*(*l*). We observed that 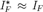 for all networks investigated in this work (Fig. 1f, inset). This result shows that for a large variety of networks structures the outdegree is the dominating factor that determines network inferability.

Inference of transcriptional networks on a genome scale is best realised by methods that are (i) asymptotically unbiased, (ii) scalable to large network sizes, (iii) sensitive to feed-forward loops^10^, and (iv) can handle data sets with and without knowledge about which nodes are targeted by experimentally induced perturbations ^7^, ^11^–^15^ (Supplementary Note 2). Inference methods for directed networks typically require individual perturbation of all nodes ^7^ or many perturbations of different strengths to compute conditional association measures ^6^,^16^ or conditional probabilities ^17^. Generation of time course data seems to be the most natural way to infer directed networks by simply analysing the temporal ordering of the transcriptional activities ^18^, ^19^. However, this approach precludes the use of knockout experiments and requires fast acting perturbations in combination with monitoring node activities over time, which is experimentally demanding, especially if nodes respond on very different time scales ^20^.

Inference is further complicated by the fact that transcriptome data contains a significant amount of stochastic variation between biological replicates despite pooling over millions of cells (Supplementary Fig. 1a). It is interesting to see that the variation across biological replicates for baker’s yeast ^3^ is close to a normal distribution and follows almost exactly a *t*-distribution with 11 degrees of freedom over five standard deviations (Supplementary Fig. 1a, inset) and that biological noise is much larger than technical noise (Supplementary Fig. 1b). As biological noise may arise from subtle differences in growth conditions that induce changes in gene regulation, we expected to see significant cross-correlations among genes (Supplementary Fig. 1c), whose magnitude is much larger than expected by chance (Supplementary Fig. 1c, inset). These cross-correlations can be exploited for inferring the structure of undirected networks ^11^, if the driving noise is independent and identically distributed for all nodes (Supplementary Note 2). In contrast, technical noise not only reduces the statistical significance for detecting true interactions but can also induce a significant fraction of false positive interactions, especially if the interaction network under investigation is sparse. Such noise induced misclassification of links can be illustrated by a simple linear network A → B → C for which standard inference methods interpret the information that A has about C erroneously as a direct link between A and C if the state of B is corrupted by measurement noise (Supplementary Fig. 1d). The reason is that a part of the correlation between A and C cannot be explained by B.

To make use of the rapidly growing amount of single-gene knockout screens for which transcriptome data is ^3^ or may become available ^21^, ^22^, we developed a method to infer directed networks on a genome scale where the number of genetic perturbations is typically below the number of nodes or genes in the network (Online Methods). In brief, our method uses the concept of probabilistic principle component analysis ^23^ to compute partial response coefficients (PRC) that are asymptotically unbiased with respect to Gaussian measurement noise (Supplementary Fig. 1d). Additionally the algorithm provides a feature to identify non-inferable links, which are removed before statistical analysis. In absence of noise, our numerical method correctly predictes the fraction of links that are inferable, *F*(*q*) (Supplementary Notes 1). To evaluate the performance of our method we generated two synthetic knockout data sets that closely resemble the gene regulatory network structure of baker’s yeast, using the GeneNetWeaver software ^24^ that uses a hierarchical network structure and our own generative model that uses a scale free network structure (Supplementary Notes 3). We added Gaussian measurement noise to the synthetic data with a standard deviation of 10% the log_2_ fold-change in expression level for each perturbation for each gene. Residual bootstrapping among replicates was used to quantify the statistical significance of the inferred link strengths. In comparison with standard inference methods, such as partial correlations ^11^–^13^, ^25^,^26^, our method shows a significantly higher performance in absence of any penalties that enforce sparse network structures (Fig. 2b, left panel). The improved performance of our method can be assigned to the fact that it is unbiased with respect to measurement noise (Online Methods).

**Figure 2.**
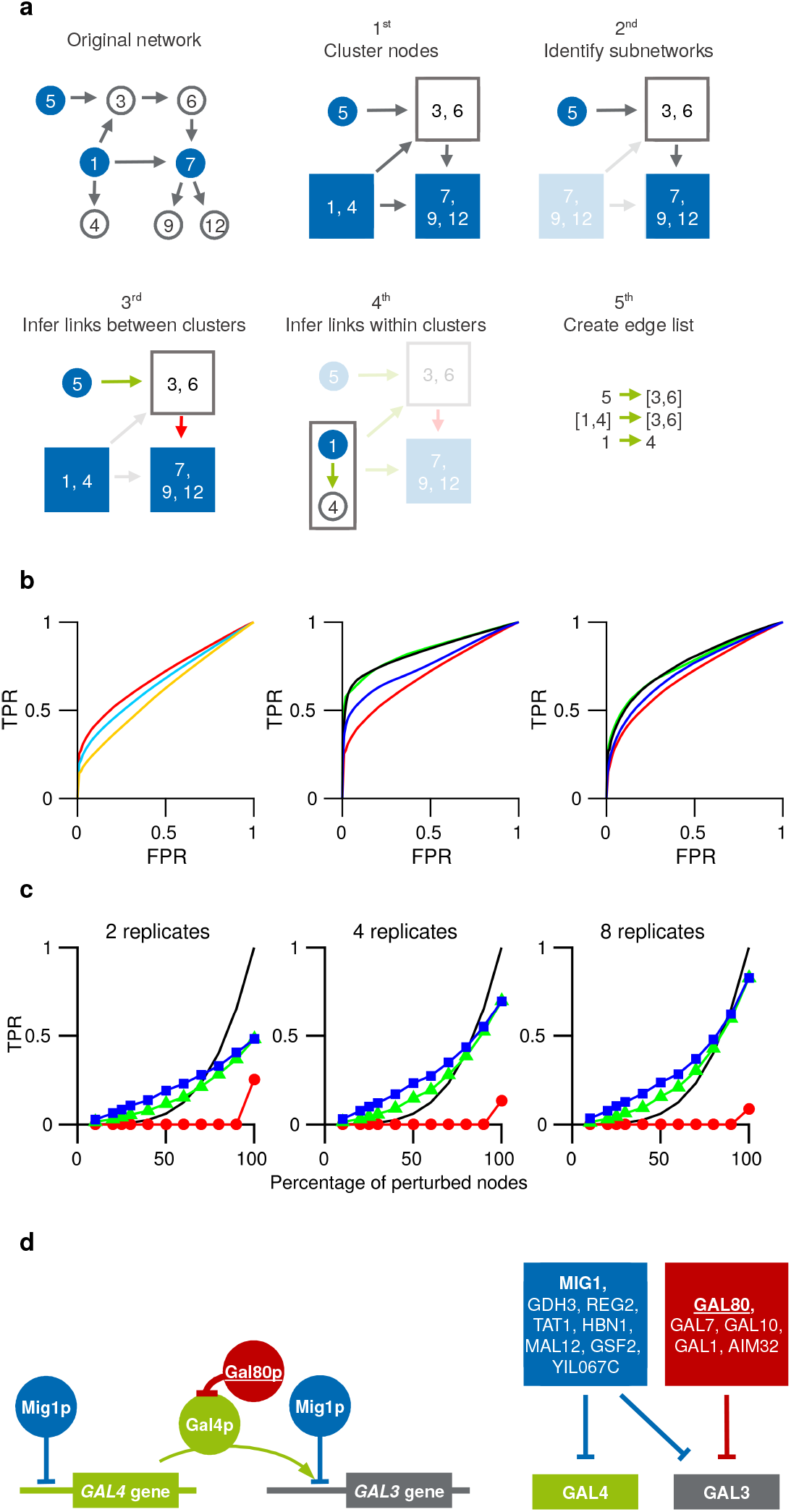
Performance of our method. **a)** Flow-chart showing the algorithmic steps for network inference as explained in the main text. **b)** Receiver Operating Characteristic (ROC) curves for 300-node scale-free networks with additive Gaussian measurement noise of 10% of the expression level and 25% of the nodes perturbed. Data was generated using GeneNetWeaver (left and middle panel) as well as using scale-free network structure with mean degree of ⟨*k*⟨ = 2 and scaling exponent *γ* = 2.5 (right panel, Supplementary Notes 3). Here, the true positive rate is computed with respect to the inferable links^30^. Performance of inference methods without sparsity constraints (left panel): PRC (red), partial correlations / linear regression (turquoise) and conditional mutual information (orange). Performance of inference methods with sparsity constraints (middle and right panel): PRC with subnetwork method (green) and Lasso (black) both applied to a subset of significantly responding nodes that were selected with 1% false discovery rate, Lasso regression applied to all 300 nodes (blue), and PRC from left panel (red) for comparison. c) True positives for the same scale-free network of (b) with 2, 4 and 8 experimental replicates with 5% false discovery rate for both significantly responding nodes and link strength: PRC (red), PRC with subnetwork method (green), PRC with subnetwork and clustering method (blue), and *F(q)* (black line). d) The *S. cerevisiae* GAL network as an example for a gene regulatory network where phosphorylated Mig1 sets the basal expression levels of Gal4 and one of its many regulatory targets Gal3. Gal4 protein can activate Gal3 but is inactivated upon binding of Gal80 protein. The transcriptome dataset contains knockout mutants for Gal80 and Mig1 but not for the remaining Gal genes. A schematic representation of the key molecular mechanisms (left) and links inferred from transcriptome data ^3^ (right).

To further improve the predictive power of our method we included the prior knowledge that transcriptional networks are highly sparse. Sparsity constraints are typically realized by penalising either the existence of links or the link strengths by adding appropriate cost functions, such as *L*^1^-norm regularized regression (Lasso) ^27^. Adding a cost function to the main objective comes with the problem to trade-off the log-likelihood against the number of links in the network whose strength is allowed to be non-zero. In absence of experimentally verified interactions there is no obvious way how to determine a suitable regularization parameter that weights the likelihood against the cost function, which is one of the great weaknesses of such methods.

In our approach we reduce network complexity by assuming that functionally relevant information in molecular networks can only pass through nodes whose response to perturbations is significantly above their biological noise level. The individual noise levels can be estimated from natural variations between wild type experimental replicates (Supplementary Fig. 1). The significance level that removes nodes from the network with high noise-to-signal ratio can be set by the desired false discovery rate. It can be shown that removal of noisy nodes imposes a sparsity constraint on the inference problem (Online Methods). The different steps required to arrive at a list of significant links are illustrated in Fig. 2a. In the first step, genes are grouped in clusters that are co-expressed under all perturbations. These clusters are treated as single network nodes in the subsequent steps. In the second step, only those samples are extracted from the dataset that correspond to perturbation of a chosen gene – the source node – with no other genes perturbed (node 5 in Fig. 2a). From this reduced dataset, we identify all nodes in the network that change expression above a given significance level upon perturbing the source node. These significantly responding nodes define a subnetwork for each source node, which is typically much smaller in size than the complete network. In the third step, we collect all perturbation data from the complete dataset for all nodes that are part of the subnetwork. Before inferring a direct interaction that points from the source node to a given target node in the subnetwork, we remove the perturbation data of target node and all nodes that respond significantly to perturbations of the target node (green arrows in Fig. 2a). The second and third steps realise numerically the counting procedure of inferable links as illustrated in Fig. 1a. Significant links are identified by partial response coefficients in combination with residual bootstrapping over replicates (Online Methods). In the fourth step, we collect all clusters of co-expressed genes that contain two nodes in total with one of the nodes perturbed and check statistical significance of the directed link between them. In the fifth step, all significant links are collected in an edge list. We refer to these five steps as the clustering method. If we remove all links from the edge list that have more than one node as a source or more than one node as a target, we obtain an edge list that corresponds to links between single genes. This reduced edge list would also arise by skipping the clustering step and we refer to remaining inference steps that compute links between single genes as subnetwork method.

The Lasso method followed by bootstrapping has been benchmarked as one of the highest performing network inference methods for in silico generated expression data ^28^. The receiver operating characteristic (ROC) curve of the subnetwork method shows better performance than the Lasso method (Fig. 2b, middle and right panel) after adjusting the regularization parameter of the Lasso method such that the area under the ROC curve is maximized. However, a significant performance boost for the Lasso method can be achieved by applying the second step of our method that removes noisy nodes, resulting in comparable performance of Lasso with the subnetwork method for the case that validation data exists such that the regularisation parameter can be determined (Fig. 2B, middle and right panel).

To get insight into the optimal experimental design for generating data for network inference, we computed the fraction of correctly inferred links and compared them against the fraction of independently perturbed nodes for different numbers of experimental replicates. We compared three different variants of our approach: partial response coefficients (PRC), PRC together with subnetwork method, PRC together with clustering method (Fig. 4C). As all variants share PRC as underlying inference method (Online Methods), the observed strong increase in performance can be assigned to the sparsity constraint that comes with the subnetwork method or the clustering method. Due to this constraint, both the subnetwork method and the clustering method can have higher accuracy than the noise-free analytical solution, as the latter does not enforce sparse network structures. The results show that in presence of 10% measurement noise the amount of available replicates limits the true positive rate, even if 100% of nodes are perturbed. However, inference of more than 80% of the network can only be achieved if the number of replicates is sufficiently high.

To evaluate the performance of our approach on real data, we use one of the largest publicly available transcriptome data sets ^3^, compromising transcriptomes that cover 6170 genes for 1441 single gene knockouts. We use the galactose utilisation network as a gene regulatory example, which is one of the best characterised gene regulatory modules in yeast ^29^. The regulatory mechanism of the GAL4 gene as a key regulator is shown in Fig. 4D, left panel. As information about phosphorylation and protein interaction is absent in expression data, the inferred network structure from transcriptome data with GAL4 and GAL80 perturbed is different from the known gene regulation. By sorting genes with respect to their number of statistically significant outgoing links, we can identify potential key regulators. Besides transcription factors, the regulators with highest statistical significance are factors involved in chromatin remodelling, signalling kinases, and genes involved in ubiquitination (Supplementary Tables 1–3). This result – although expected for eukaryotes – is inaccessible for inference methods that a priori fix known transcription factors as regulatory sources. However, as the 1441 knockout genes of this dataset compromise just 23% of the genes for which transcript levels have been measured, we can estimate from our simulations that we have inferred less than 10% of the direct interactions in the transcriptional network of yeast. However, the recently available double knockout screens in yeast ^22^ drastically increased the number of targeted perturbations, which can boost the prediction accuracy of network inference methods once transcriptome data for a fraction of these mutants become available.

**Supplementary Figure 1.**
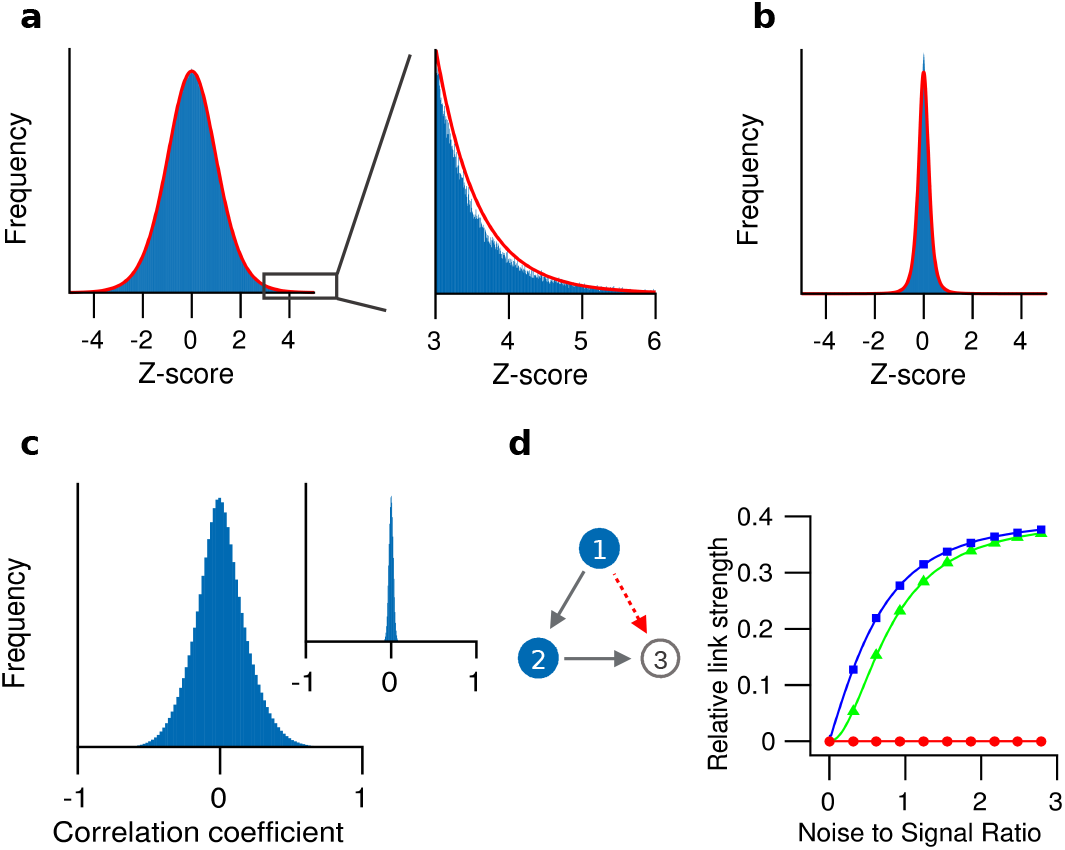
Distribution of wild type expression levels for *S. cerevisiae* from 748 biological replicates. **(a)** Distribution of the relative expression, *log*_2_(*r*_*i*_), with 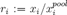 and *x*_*i*_ the expression of gene i relative to 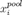 followed by standardization of the log_2_ fold changes (z-score). The values 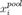 have been obtained by first pooling the 748 biological replicates before measuring gene expression. The distribution is well described within 5 standard deviations by a t-distribution with 11 degrees of freedom (red line). **b)** Distribution as in (a) but now for differences among technical replicates. **c)** Correlation between gene expression levels is significantly higher than expected by chance (inset) **d)** Illustration of a noise induced false positive link (red arrow) as described in the main text. Data for a three-node network with two links was generated by applying independent perturbations on node 1 and node 2. The link strength of the non-existing link from node 1 to node 3 relative to the existing link from node 2 to node 3 was inferred using three different methods (i) partial correlations (blue squares), (ii) conditional mutual information (green triangles), and PRC (red circles).

## Methods

### Partial response coefficients (PRC)

We aim at inferring direct interactions between *N* observable molecular components, such as transcripts or proteins, by measuring their copy numbers or concentrations, *y ϵ* ℝ^*N*^. We assume that the available dataset has been generated from *P* perturbation experiments, 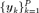, which may also include experimental replicates. We further assume that the molecular targets of the perturbations are known, as it is the case for gene knockout or knockdown experiments. The elements of the interaction matrix *A* ϵ ℝ^*NxN*^ define the strength of the directed interactions among the molecular components, for example *A*_*ij*_ quantifies the direct impact of component *j* on component *i*. Given the available experimental data, our aim is to correctly classify the off-diagonal elements of ***A*** as zero or nonzero to obtain the structural organization of the interaction network. We assume that the observed component abundance, 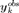, differs from the true value, ***y***_*k*_, by additive measurement noise, *ϵ*(*t*), which is characterized by zero mean, 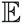[*ϵ*(*t*)] = 0, and variance, 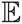[*ϵ*(*t*)*ϵ*(*t*)^*T*^] = *σ*^2^*IN*, with *I*_*N*_ the *N* dimensional identity matrix. We assume that the observed data can by described to sufficient accuracy by a linear stochastic process

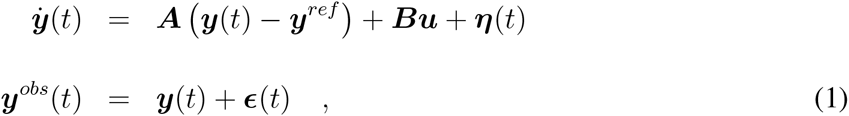

with ***A*** negative definite to ensure stability. Equations of this type typically arise from linear expansion of a nonlinear model around a reference state, *y*^*ref*^. The perturbation vector ***u*** is time-independent and reflects perturbations that persist long enough to propagate through the network, such as mutations that affect gene activity. We further introduce with *B* ϵ ℝ^*NxN*^, the associated standard deviations of the perturbative forces, ***u***, and assume that these forces are sampled from a standard normal distribution, with mean 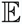[*u*] = 0 and covariance matrix 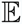[*uu*^T^] = *I*_*n*_. In general, only the positions of the non-zero elements of *B* are known from the experimental setup but their actual values are unknown. Fluctuating perturbations that are fast in comparison to the network response are represented by the stochastic vector *η*(*t*), which we model as white noise defined by 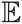[*η*(*t*)] = 0 and 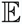[*η*(*t*)*η*(*t′*)] = *γδ*(*t – t′*) *I*_*N*_, with *δ*(*t*) the delta function and *γ* the fluctuation strength. The solution of Eq. (1) is given by

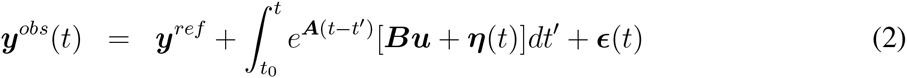

from which we obtain in the stationary regime, *t ≫t*_0_, a relation between ***A*** and the covariance matrix of observed component abundances

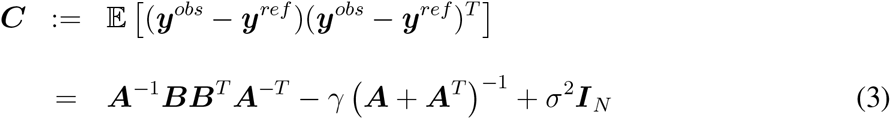

We exploit Eq. (3) to infer directed networks from correlation data. Here, we assume that component abundances are obtained from averaging over a large number of cells in the stationary growth phase. In this case, fast fluctuating perturbations that arise from thermal noise and can be observed only on single-cell level average out. To infer the interaction matrix, ***A***, we start with singular value decomposition of the matrix product *A*^-1^*B*

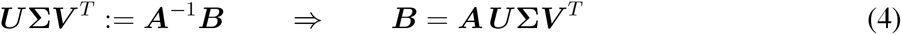

with ***U*** and ***V*** orthogonal matrices and ∑ a diagonal matrix containing the singular values. The negative definite matrix ***A*** is invertible and hence has full rank. In the following, we show that it is possible to infer the directed link strength between a sender node *j* and a receiver node *i*, if all direct perturbations on receiver node *i* are removed from the dataset and if a significant partial correlation between *i* and *j* exists. Removing the perturbation data for node *i* implies that the matrix *B* has at least one zero entry. As a consequence, *N*_0_ ≥ 1 singular values are zero – as in general not all nodes are perturbed – and the corresponding rows of *U* span the left nullspace of ***A***^−1^***B***. In the absence of fast fluctuating perturbations, *γ* = 0, we can rewrite the covariance matrix as

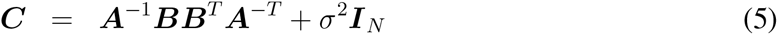

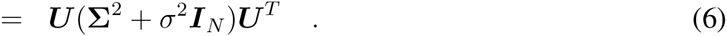

Assuming that the observed node activities follow a multivariate normal distribution, we can find estimates for the unknown orthogonal matrix *U*, the singular values ∑, and the observational noise *σ* by maximizing the log-likelihood function 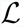 under the constraint 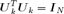, with

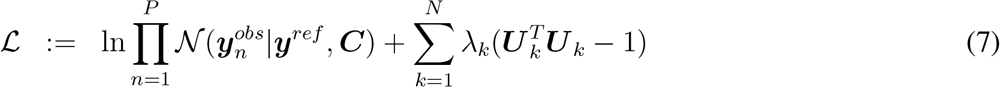

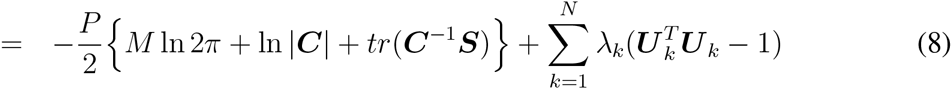

Here, 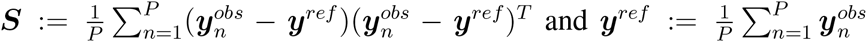 denote maximum likelihood estimates of the covariance matrix ^31^ and the expected node activities, *λ*_*k*_ is a Lagrange multiplier and *tr*(·) denotes the trace of a matrix. In the following calculations, we substitute *S* by the unbiased sample covariance matrix, *S*→ *P*(*P* — 1)^−1^*S*. Note that *V* must disappear in the likelihood function as the covariance matrix of *u* is invariant under any orthogonal transformation *u* → *V*^*T*^*u*.

The maximum of the log-likelihood function is determined by the conditions *d*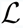/*dU*_*k*_ = 0, *d*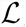/*d∑*_*kk*_ = 0, and *dL*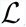/*dσ*^2^ = 0, which results in

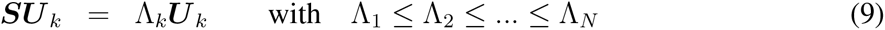

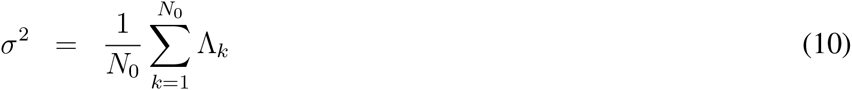

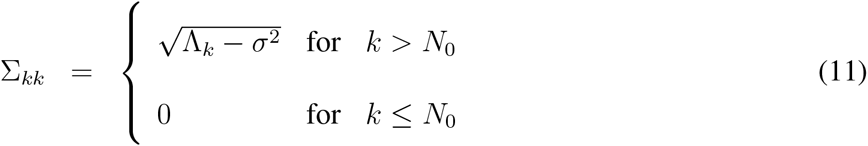

showing that maximum likelihood estimates of *U*, *σ*^2^, and ∑ are uniquely determined by the sample covariance matrix *S*. Our analysis is mathematically equivalent to the probabilistic interpretation of principle component analysis ^23^.

Solving the matrix equation, Eq. (4), for *A* gives

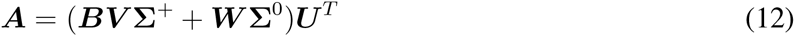

with ∑^+^ the pseudoinverse of ∑. As the matrix *A* has full rank, we complement ∑^+^ with an unknown diagonal matrix ∑^0^ that has nonzero values where ∑^+^ has zero values and vice versa and complement *BV* with an unknown orthogonal matrix *W*. Note that by construction, *∑*^+^*U*^*T*^ and *∑*^+^*U*^*T*^ map from complementary subspaces and thereby ensure that *A* has full rank. The fact that *V, W*, and ∑^0^ cannot be determined from *S* shows that *A* cannot be computed from a single covariance matrix. A more general case arises when measurement noise is independent but not isotropic, *σ*2*I* → *σ*^2^*D*, with *D = diag*(*r*_1_, *r*_2_,…, *r*_*N*_) a diagonal matrix with known positive elements that contains scaled noise variances, 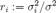, resulting in

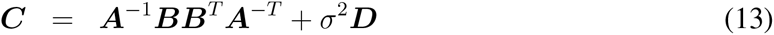

A transformation to isotropic noise is possible by multiplying both sides of Eq. (13) by 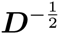, which changes the result Eq. (12) to

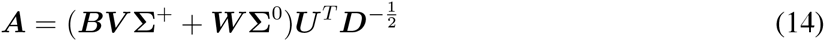

with *U* the eigenvectors of 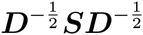.

### Case *N*_0_ = 1

We assume that the i-th node is the only unperturbed node in the network and consequently define *B*_*il*_ = 0 for all *l*. From Eq. (12) we obtain a unique solution for the *i*-th row of *A* relative to the diagonal element, *A*_*ii*_,

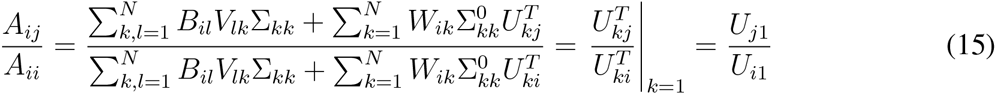

with *U*_*j*1_ the *j*-th element of the eigenvector of *S* that has the smallest eigenvalue. Note that the first term in the brackets vanishes as *B*_*il*_ = 0 and 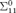 is the only nonzero element of ∑^0^. The important point is that any dependency on *σ* has dropped out, which makes this method asymptotically unbiased with respect to measurement noise. The fact that we can determine the elements of the *i-th* row of *A* only relative to a reference value, *A*_*ii*_, is rooted in fact that we have to determine the *N* parameters {*A*_*il*_,…, *A*_*ii*_,…, *A*_*iN*_} from *N* – 1 perturbations. As a consequence, the strengths of the links onto the target nodes cannot be compared directly if their restoring forces or degradation rates, *A*_*ii*_, are different. Generally, only relative values of *A* can be determined, as the average perturbation strength on node *i* cannot be disentangled from its restoring force *A*_*ii*_ – a problem that is typically circumvented by defining *A*_*ii*_ := 1 for all *i* ^7^,^12^,^14^. For the case that all nodes in the network are perturbed one-by-one, we can cycle through the network and remove the perturbations that act on the current receiver node, while keeping the perturbations on the remaining nodes. By computing the *N* corresponding covariance matrices and their eigenvectors, we can infer the complete network structure from Eq. (15) if the data quality is sufficiently high.

### Case *N*_0_ > 1

For all *i* that are not perturbed or whose perturbation data has to be removed from the dataset we get from Eq. (12)

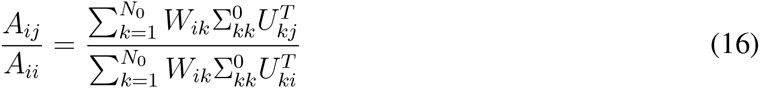

Non-unique solutions of Eq. (16) can arise if a given fraction of the variance of the receiver node *i* can be explained by more than one sender node, for example, when a perturbed node *j* targets two nodes with index *i* and *l*. In this case it is unclear from the node activity data whether *i* is affected directly by *j* or indirectly through *l*, or by a combination of both routes. If the node *l* is unperturbed, we can use the simple criteria shown in Fig. 1a to decide whether the link from *j* to *i* is inferable or not. If node *l* is weakly perturbed, a statistical criteria is needed to decide about inferability, which can be computed numerically as follows: To find out whether *j* transmits a significant amount of information to *i* that is not in *l*, we first remove the perturbed node *j* from the network and determine the link strengths *A′* for the remaining network of size *N* — 1. To construct a possible realisation of *A′* we set in Eq. (16) the nonzero values of Σ^0^ to unity and use *W* = *U* to arrive at the expression

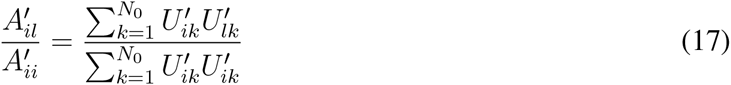

with *U′* determined from the sample covariance matrix with the *i*-th column and *i*-th row removed. Using the inferred link strength from Eq. (17) we can rewrite Eq. (3) as a two-node residual inference problem between *j* and *i*, where we obtain a lower bound for link strength from node *j* to *i* by using the variation of *i* that could not be explained by *A′*. Defining by *A͂*, *B͂* and *D͂* the 2×2 analogs to the full problem we obtain

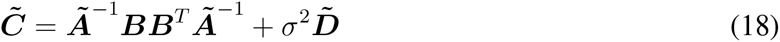

with 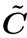 the covariance matrix of the vector 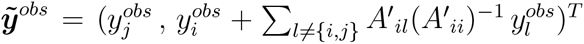 and 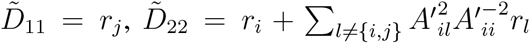, using the scaled variances 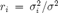. Note that *A*_*ii*_ < 0 for all *i* as these elements represent sufficiently strong restoring forces that ensure negative definiteness of *A* and that we have 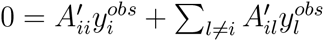 from Eq. (1) in the stationary case. An estimate for the minimum relative link strength from node *j* to node *i* can be calculated from Eq. (14) and is given by

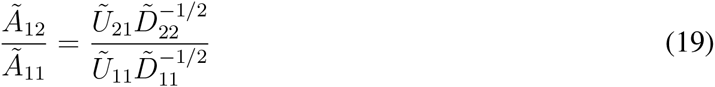

Eq. (19) can be considered as an asymptotically unbiased response coefficient between node 1 as target node and node 2 as source node, as any dependency on *σ*^2^ has dropped out. An estimate for the maximum relative link strength from node *j* to node *i* follows from Eq. (19) with the off-diagonal elements of *A′* set to zero. We classify a link as non-inferable if there exists (i) a significant difference between the minimum und maximum estimated link strength and (ii) a minimum link strength that is not significantly different from noise.

### Computational complexity of PRC

The computational cost for computing partial response coefficients scales as 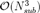, with *N*_*sub*_ the size of the subnetwork under consideration. However, as we infer directed networks, we first have to remove the perturbations on each target node before its incoming links can be inferred. The cycling through up to *N*_*sub*_ — 1 perturbed target nodes increases the computational complexity to 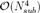 in the worst case. As we have generated a subnetwork for each significantly varying node and used residual bootstrapping to infer statistically significant links, the total computational complexity is given by 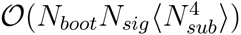, where 〈·〉 denotes averaging over all subnetworks and *N*_*boot*_ the number of bootstrap samples. If the travelling distance of perturbations (correlation length) in the network is significantly shorter than the network diameter, such that *N*_*sub*_/*N* → 0 in the limit of large networks, *N* → ∞, the computational complexity scales linear with network size. In contrast, using Lasso to infer directed links requires 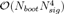 operations, as the more efficient Graphical Lasso method ^32^ is only applicable to undirected networks. Whether our method is computationally more efficient than Lasso depends on the inference problem. However, for the scale free networks investigated in this work our method required significantly less computational time than inference via Lasso using parallel computing.

### Fraction of non-inferable links

Inferability of a directed link between source and target node requires that the remaining network may not contain the same information that is transmitted between them. A sufficient condition is that all information that the remaining network receives from the source node is destroyed by sufficiently strong perturbations. If the target node is not perturbed, information from the source node my reach the remaining network through the target node. In this case also the targets of the target node must be perturbed (Fig. 1a). Counting network motifs that satisfy these conditions gives the number of inferable links. If the network size, *N*, is significantly larger than the number of outgoing links for both the source and target nodes, we can approximate the fraction of inferable links, *F*(*q*), by the expression (Supplementary Note 1)

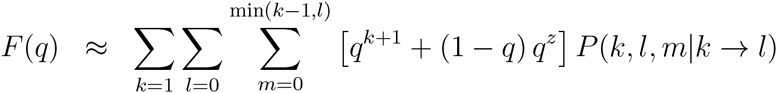

Here, *P*(*k*,*l*,*m*|*k* → l) is the conditional probability of finding two connected nodes in the directed network, where the source node has *k* ≥ 1 outgoing links, the target node has *l* ≥ 0 outgoing links, and both share *m* nodes as common targets of their outgoing links. The first term in the brackets corresponds to the case that independent perturbation data for node B exists (Fig. 1a, left panel) and the second term to the case where independent perturbation data for node B is absent (Fig. 1a, right panel). In the calculation of *F*(*q*) we assumed that the links in the network are identical with respect to the information they can transmit.

### Data preparation

Kemmeren et al.^3^ provided a transcriptome data set of a Saccharomyces cerevisiae genome-wide knockout library (with mutant strains isogenic to S288c). This data set comprises transcript levels of 6170 genes for 1484 deletion mutants. The data is presented as the logarithm of the fluorescence intensity ratios (M-values) of transcripts relative to their average abundance across a large number of wild type replicates, resulting in logarithmic fold changes of mutant/wild type gene expression levels compared to a wild type reference level. Kemmeren et al. also used a dye swap setup for several experiments to average out the effect of a possible dye bias. Their chip design measures most of the genes twice per biological sample, thus allowing to estimate the technical variance. The preprocessing of the data is described in Kemmeren et al.^3^, Supplementary Information.

### Residual Bootstrapping

We make use of the 748 measured wild type experimental replicates to determine the natural variation among biological replicates, 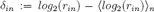, with 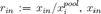 the expression of gene *i* in wild type replicate 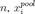 the expression level of gene *i* measured after pooling over wild type replicates, and 〈.〉_*n*_ denoting the average over replicates. To generate the bootstrap samples we randomly select 200 different *δ*_*in*_ from the replicates for each gene *i*, and add these values to the log fold-changes of the perturbed expression levels, 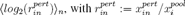 and the average is taken over the two replicates for each knockout.

### Sparsity constraints by removing noisy nodes

As network inference typically comes with an insufficient amount of independent perturbations and experimental replicates we run into the problem of overfitting the data. In this case, noisy information from many network nodes is collected to explain the response of a given target node. *L*^1^-norm regularized regression (Lasso) systematically removes many links, where each link explains only a small part of the variation of the target node, in favour of few links, where each link contributes significantly. In our approach we remove noisy nodes and thus their potential outgoing links, where the critical noise level is determined by independent biological noise. Which nodes are removed depends on the *source node* that is perturbed. In the absence of noise, our algorithm removes weakly responding nodes from the network. We thereby assume that the existence of many indirect interactions between source and target node by first distributing the signal of the source node among many weakly responding nodes and then collecting these week signals to generate a significantly responding target node is much less likely than the existence of a single direct interaction. However, in the noise-free case we run into the same problem as Lasso to determine the right cut-off (regularization parameter).

### Synthetic Data

Synthetic data was generated using our own model and GeneNetWeaver ^24^ – an open access software that has been designed for benchmarking network inference methods. With GeneNetWeaver, networks were generated from a model that closely resembles the structure of the yeast regulatory network ^24^, and steady state levels of node activities were computed using ordinary differential equations (ODEs). In our model, we first generated scale-free networks with an exponent of 2.5 and an average degree of 2. Then, we solved a system of ordinary differential equations with non-linear regulatory interactions between nodes to obtain steady state values of node activities, e.g. transcript levels. For both models, logarithmic fold changes of node activities were calculated (transcriptional levels upon perturbation relative to wild levels), and gaussian measurement noise was added.

## Acknowledgements

We acknowledge help from Patrick Kemmeren in interpreting the yeast transcriptome data, the High Performance Computing Plattform of the Heinrich-Heine University, and DFG funding by SPP 1395 and the Cluster of Excellence in Plant Sciences (grant no. EXC 1028).

## Author contributions statements

A.S.K. and M.K. conceived the PRC method. N.H. and M.K. developed the counting procedure for inferable links and the Inferability measure. C.F.B. and M.K. developed the clustering and subnetwork methods and analysed the transcriptome data. N.H. wrote Supplementary Note 1. M.K. wrote Supplementary Note 2. C.F.B. wrote Supplementary Note 3.

## Correspondence

Correspondence and requests for materials should be addressed to M.K.

## Supplementary Note 1

### 1 Counting the number of inferable links with respect to the number of perturbed nodes

Structural features of a network allows us to determine the expected fraction of inferable links analytically if the number of randomly perturbed nodes (*N*_*p*_) is known and measurement noise is absent.

Assume a subnetwork where a source node (A) targets a node (B), given that the out-degree of node A is *k*, the out-degree of node B is *l*, and A and B have *c* nodes as common targets. We aim to infer the link from node A to node B (See figure 1).

**Fig. 1.**
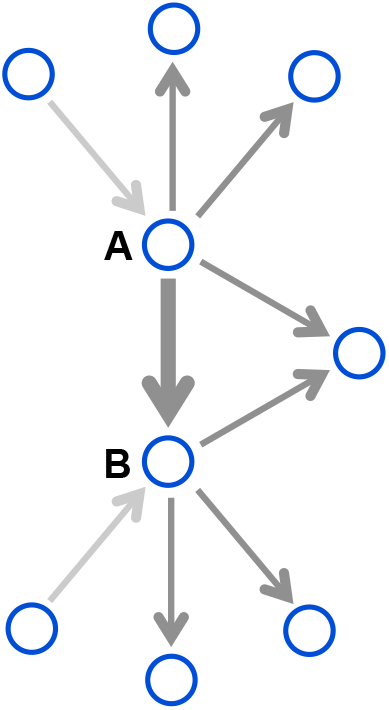
Example of a directed subnetwork. We would like to infer the link from node A to node B (shown in bold), where A has out-degree 4, B has out-degree 3, and A and B share 1 common target.

We denote a direct link from source node (A) to target node (B) as inferable if there exits a detectable amount of mutual information between A and B that cannot be transmitted by any alternative route through the network. This requires that at least one nodes of each alternative route is perturbed and thereby part of the transmitted information is destroyed. To meet this requirement, either one of the following conditions must be fulfilled (Fig. 1a, main text):

1. All nodes that are targeted by A are perturbed, including node B.
2. Except B, all nodes that are targeted by A and B are perturbed.

By collecting all subnetworks that fulfil conditions 1 or 2 we can calculate the average fraction of inferable links, *F*(*N*_*p*_), using *N′* := *N*–*k*–1 and 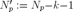

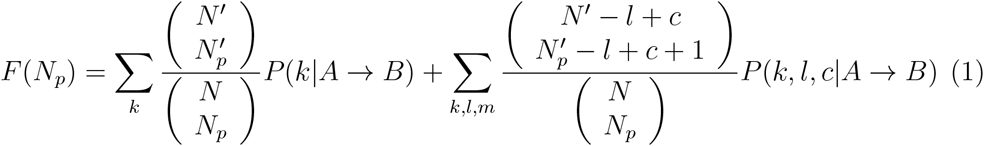

Here, *N* is network size, *N*_*p*_ is number of perturbed nodes, *k* is the out-degree of the source node (A), *l* is the out-degree of the target node (B), *c* is the number of common nodes targeted by A and B. We further defined by *P*(*k*|*A* → *B*) the conditional probability that for any two connected nodes, source node (A) has out-degree *k* and *P*(*k*, *l*, *c*|*A* → *B*) is conditional probability that for any two connected nodes, source node (A) has out-degree *k*, target node (B) has out-degree *l*, and A and B target *c* common nodes. The first term in the numerator counts motifs that fulfil condition 1, and the second term in the numerator counts motifs that fulfil condition 2. The term in the denominator counts all possible network motifs, when *N*_*p*_ nodes of the network are perturbed. Note that measurement noise can decrease the fraction of inferable links due to distortion of information flow. Thus equation 1 calculates the expected maximum fraction of inferable links.

Assuming that the network size, *N*, and the number of perturbed nodes, *N*_*p*_, are much larger than the out-degrees, *N*, *N*_*p*_ ≫ *k*,*l*, we can apply Stirling’s approximation and simplify Eq. (1)

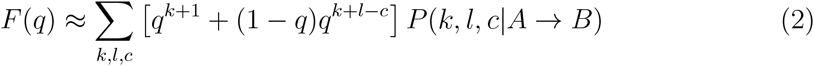

where *q* denotes fraction of perturbed nodes, *q* = *N*_*p*_/*N*.

### 2 Network’s Inferability

According to Eq. (2), networks with different structural features have different *F*(*q*) curves (Fig. 1b, main text). We therefore define a inferability measure, *I*_*F*_, as the area under the curve of F(q) that reflects how difficult it is to infer links for a given network structure

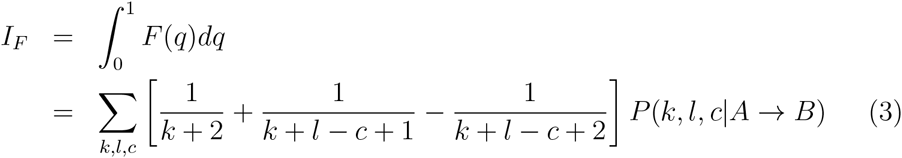

Not that *I*_*F*_ is independent of network size. For a sufficient large networks *N* ≫ 1 and when feed forward loops rare, we can approximate the joint probability by

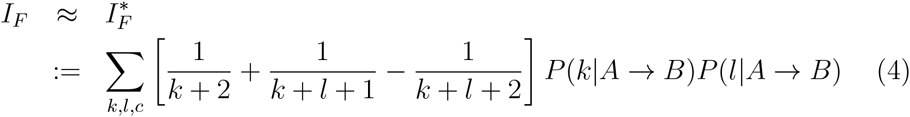

According to Eqs. (3) and (4), inferability mainly depends on the out-degree of nodes. Consequently, networks consisting of nodes with high out-degree have low inferability (*I*_*F*_) and are the most difficult ones to infer.

### 3 Comparison of *F*(*N*_*p*_) with the inference algorithm

We compared the fraction of inferable links determined by the analytical formula, *F*(*N*_*p*_), and the average number of inferable links classified by our inference algorithm in the limit of low measurement noise. In order to calculate the average number of inferable links form the inference algorithm, first we consider a specific network type. For each number of perturbed nodes, *N*_*p*_, we first randomly identify nodes that will be perturbed and subsequently generate node activity data. For each random sample of *N*_*p*_ nodes, we calculate the fraction of inferable links and finally average over all configurations of perturbed nodes. Figure 2 compares the results of the analytical and numerical approaches for three different types of networks. We consider the two extreme examples of scale-free networks; a scale-free network where hubs are targets of links (Fig. 2a), a scale-free network where hubs are souces of links (Fig. 2b) and a random network (Fig. 2c). In all three network types the network sizes are *N* = 100 and averaging runs over 300 different configurations of *N*_*p*_ randomly selected nodes that are perturbed individually by knockouts.

**Fig. 2.**
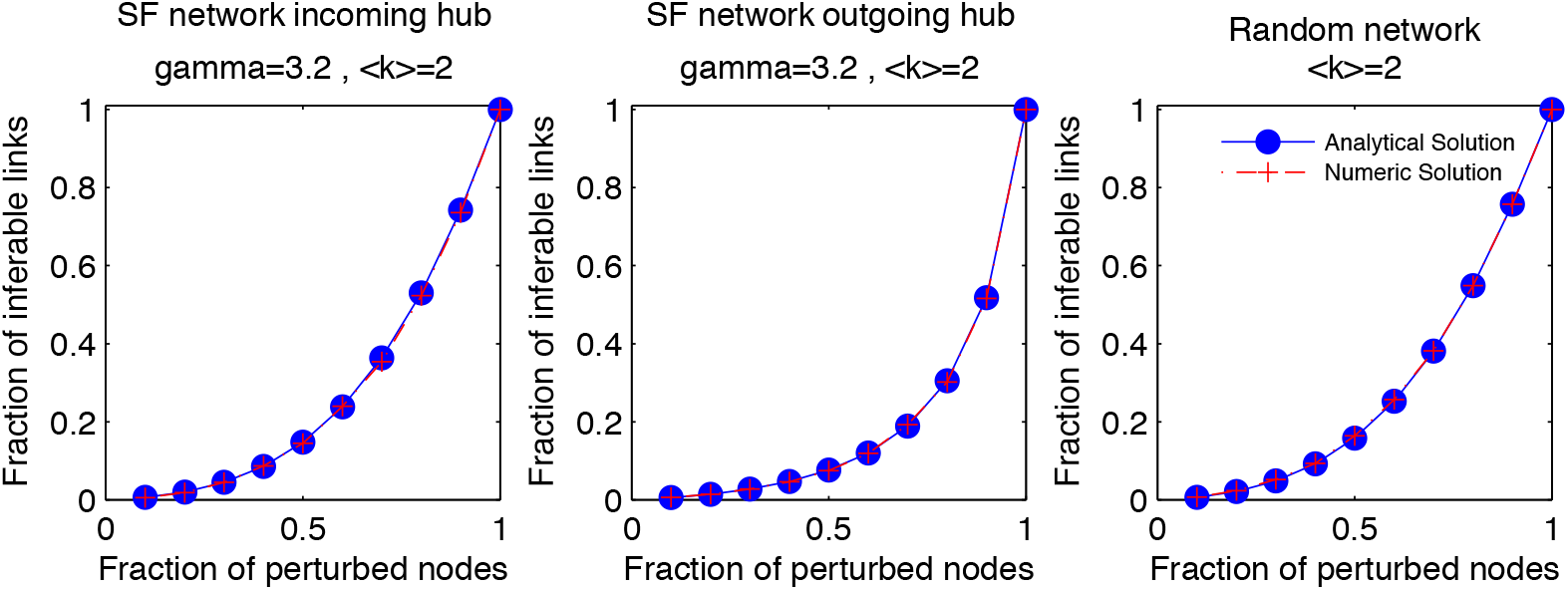
Comparison of the analytical formula, Eq. (1) and the inference algorithm by calculating the number of inferable links with respect to the number of perturbed nodes, *N*_*p*_.

## Supplementary Notes 2

### Interpretation of network inference algorithms that are related to partial correlations

Algorithms based on partial correlations [1, 2] – a method designed to discriminate between direct and indirect interactions – show comparatively low performance for predicting direct correlations between transcript levels using transcriptome data [3] but excellent performance for predicting contact points of protein structures [4, 1]. To understand the difference in performance of partial correlations, we investigate Eq. (3) of the Online Methods section for the special case where (i) all perturbations are independent and fast fluctuating, B = 0, (ii) all links are bidirectional (undirected network), *A* = *A*^*T*^, and (iii) measurement noise is absent, *σ* = 0. In this case the elements of interaction matrix, *A*, correspond to partial correlations for *γ* = 2

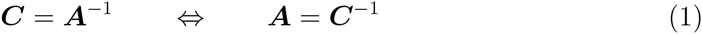

The constraints imposed by the conditions (i)-(iii) explain the poor performance of partial correlations in inferring undirected links for gene regulatory networks [3], where most of data is generated by perturbations that persist on long time scales, measurement noise is significant, and the network is directed. In contrast, conditions (i)-(iii) are satisfied to very good approximation for identifying physical contacts between amino acids in folded proteins [4], where single nucleotide substitutions occur on much shorter time scales than significant changes in protein structure, links are undirected, and measurement noise from sequencing errors make no relevant contribution.

In contrast to undirected networks, directed links can be inferred in absence of measurement noise using a response matrix method, if static perturbations are applied individually to all nodes in the network [5]. Given that fast fluctuating perturbations are absent, we can calculate from Eq. (2) of the Online Methods section the response of the *i*-th node, *δy*_*i*_, with respect to a perturbation that acts exclusively on the j-th node by 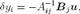, with 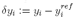 and *B*_*j*_ the *j*-th row of *B*. Using the expression 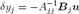 to substitute *Bju*, we arrive at the relation

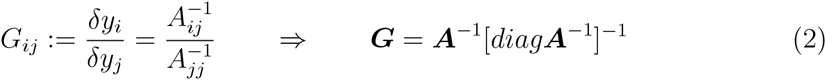

with diag ***X*** a matrix with elements {*X*_11_, X_22_, …} on the main diagonal and zero elements otherwise. Here, *G*_*ij*_ denotes the linear response coefficient of the network, describing the change in activity of node *i* if node *j* changes its value in response to a perturbation that acts exclusively on node *j*. After inverting both sides of Eq. (2) and setting the restoring forces (degradation rates) to unity, *A*_*jj*_ = –1 for all *j*, we arrive at the published result of Ref. [5]

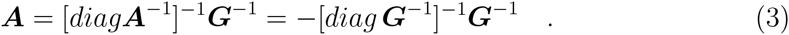

In the derivation we used that by definition *diag****A*** =–*I* = [*diagA*^−1^]^−1^ *diagG*^−1^, with ***I*** the identity matrix.

The lack of a unique relation between ***G*** and ***A*** is rooted in the fact that the *N*(*N* +1) parameters of ***A*** and ***Bu*** cannot be uniquely identified from the *N × N* measured node activities, even for the rare case that it was possible to perturb all nodes in the network one-by-one. This problem does not change if more perturbations are carried out as ***Bu*** takes different values for different perturbations. The result that direct interactions can be computed by inverting the linear response matrix, Eq. (3), has been derived several times, with either fixing the restoring force, ***A*** = –***I***, [5, 2] or by computing direct interactions from ***A*** = ***I*** – ***G***^-1^, [1] (the latter reference defines an unusual response matrix with zero elements on the main diagonal by ***G***_*obs*_ := ***G–I***).

Substantial confusion arrises if ***G*** is substituted by ***C*** in Eq. (3) for the reason that ***C*** can be computed using perturbative forces that act simultaneously on many nodes [2, 1], whereas computation of ***G*** requires that all nodes are perturbed individually [5]. Despite the fact that Eq. (1) and Eq. (3) look similar, the invalid operation ***G*** → ***C*** implies that we apply Eq. (1), which only holds for fast fluctuating perturbations, to static perturbations for which Eq. (3) has been derived. As expected, this invalid substitution gives poor performance if a substantial fraction of the dataset comes from slowly varying perturbations [3] and good performance if the conditions (i)-(iii) apply [1].

## Supplementary Note 3

### 1 Input data to network inference algorithm

#### 1.1 Saccharomyces cerevisiae genome-wide knockout library

##### 1.1.1 Description of the data set

Kemmeren et al. [1] provided a transcriptome data set of a *Saccharomyces cerevisiae* genome-wide knockout library (with mutant strains isogenic to S288c [2]). This data set, hereafter simply referred to as the yeast deletome data, comprises of transcript levels of 6170 genes for 1484 deletion mutants (hereafter referred to as perturbation experiments). The data of 1441 of these perturbation experiments can be used for network inference as the transcript levels of the deleted genes was measured with the microarray chip design. The data is presented as the logarithm of the fluorescence intensity ratios of red and green labelled microarray targets (M-values): 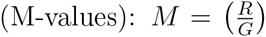. For the regular experimental setup, DNA from the deletion mutants was labelled red, and DNA that was pooled from several wild type batches was labelled green. The aim of pooling the wild type DNA was to average out biological fluctuations, so that this DNA pool could be used to define a reference gene expression level. This allows the interpretation of the M-values as logarithmic fold changes of mutant gene expression levels compared to a wild type reference. Kemmeren et al. also used a dye swap setup for several experiments to average out the effect of a possible dye bias. The chip design used by Kemmeren et al. measures each gene twice per biological sample, thus allowing estimation of the technical variance.

##### 1.1.2 Preprocessing of data

We used the preprocessed data provided by Kemmeren et al. for our analyses. The preprocessing steps are described elsewhere [1]. We re-arranged the original pre-processed data layout to make it compatible with our network inference algorithm. Specifically, M-values corresponding to dye-swap experiments were multiplied by −1 and only knock-out experiments corresponding to genes that were also included in the chip design were kept in the data set.

##### 1.1.3 Distribution of biological and technical variance (suppl. figure 1a and suppl. figure 1b)

Since there is insufficient information about the biological processes leading to variation among biological replicates (biological noise), we assume that the process is equal for all genes. By bringing the wild type data to unit variance for each gene, we show that the variation among the wild type data logarithmic fold changes is approximately Gaussian distributed (suppl. figure 1a). By taking the mean logarithmic fold change of each biological replicate (wild type data only) and then calculating the differences of the technical replicates to this mean, the distribution of the technical variance can be found (suppl. figure 1b).

##### 1.1.4 Correlation among wild type experiment M-values (suppl. figure 1c)

The histograms shown in suppl. figure 1c were created from the upper triangular correlation matrix (without diagonal elements) of wild type experiment M-values. The inserted figure was created by breaking up correlations between genes via random shuffling of the experiments.

#### 1.2 Simulating Gene Expression data

To estimate how well network inference works on data which is similar to the data set provided by Kemmeren et al. [1], the input data to the inference algorithms was generated as described below. In short, steady state solutions of ordinary differential equations were used to generate absolute gene expression levels, which were then turned into logarithmic fold changes, and measurement noise was added.

##### 1.2.1 Using scale free networks and steady state solutions of ordinary differential equations to simulate absolute gene expression levels

We chose scale free network models with exponent 2.5 to simulate a comparable network structure to yeast; the average degree was set to 2. Up to a rounding error, 80% of links were set as inhibiting. Non-linear regulatory interactions were simulated by solving ordinary differential equations. First, wild type expression levels were calculated by solving a system of equations of the form shown below. The Hill-coefficients *h*_*ki*_ were sampled from a uniform distribution between 1 and 2. The constant *K*_*ki*_ was set to 0.5. The linear degradation term *λ*_*i*_ was set to 2 to assure stability. The coefficients *a*_*ki*_ and *b*_*ki*_ were set to 1 and −1 respectively for inhibiting links and to 0 and 1 respectively for activating links.

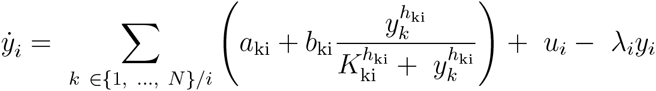

Here, *u*_*i*_ denotes the basal gene expression rate. To compute the response *y*_*ij*_ of a gene *i* to a knock-out of gene *j*, the above shown system of ODEs was modified by setting the basal expression rate for the knocked out gene to zero and by removing all links onto and away from the gene *j*.

##### 1.2.2 Using GeneNetWeaver to simulate absolute gene expression data

For better comparability with other publications, gene expression data was also generated with the program “GeneNetWeaver” [3]. This data was only used for comparing inference methods using ROC curves. From the “gold standard” yeast network in the program, random subnetworks were extracted with at least 100 regulators (random vertex seed; greedy neighbor selection). Then, the “Generate Kinetic Model” option was used with removal of auto-regulatory interactions to allow the generation of data sets. Data sets were generated with the following options: deterministic (ODEs) model, knock-out & wild type experiments, no time series, no noise added.

##### 1.2.3 Logarithmic fold changes (M-values)

To simulate logarithmic fold changes that were similar to the M-values (logarithmic fold changes) of the yeast deletome data set described above, the M-values of yeast knock-out mutants were investigated for the knocked-out genes. Due to measurement noise, these values never reach negative infinity as would be expected in the absence of technical noise. The median minimal absolute fluorescence intensity averaged over all knocked out genes was estimated to be 2^−2^.^5^. This value was added to all simulated absolute gene expression level values before calculating the logarithmic fold changes. That is, the gene expression response of a gene *i* towards a perturbation of a gene *j*, expressed as logarithmic fold change, was calculated as 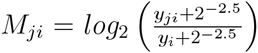, with *y*_*j*_*i* the absolute expression level of gene *i* when gene *j* is perturbed, and *y*_*i*_ the expression level of gene *i* for the non-perturbed (wild type) state, as defined above. Replicates were produced through duplication of the logarithmic fold changes for each perturbation experiment.

##### 1.2.4 Simulating measurement noise

Although the reference node activities (wild type data logarithmic fold changes) in the data set provided by Kemmeren et al. [1] are correlated, we did not simulate correlated measurement node activities. This is because we were lacking information about the processes that generate biological variance as well as the actual network structure of *S. cerevisiae*. Because the biological noise dominated the technical noise, we did not distinguish both noise types in our simulations. Rather, we simulated noise by adding normal random variables to all simulated logarithmic fold changes.

##### 1.2.5 Signal-to-noise ratio

The signal-to-noise ratio was calculated from the covariance matrix *C* by eigenvalue decomposition and analysis of the sorted eigenvalues. That is, the covariance matrix was decomposed into *C = U*Σ*U*^*T*^, where Σ is a diagonal matrix that has its eigenvalues sorted in ascending order (*σ*_11_ <…< σ_NN_) on its diagonal. Then, the signal-to-noise ratio (SNR) was calculated as 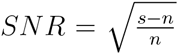, with the signal 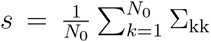 and the noise 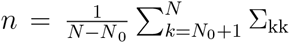. Here, *N*_0_ denotes the number of perturbed nodes, and *N* is the total number of nodes.

### 2 Workflow

#### 2.1 Data normalization

All node activities *x* were centered to the average of the corresponding reference node activities (wild type data logarithmic fold changes) *x̄*, such that the average reference node activities became zero. That is, with data for N reference node activities and *P* perturbation node activities, for a gene *i*, we have for an arbitrary normalized node activity: 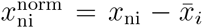, with *n* between 1 and *N* + *P* and 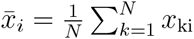.

#### 2.2 Identification of significantly affected nodes

To identify which nodes were significantly affected by a perturbation, it was checked whether the node activities were significantly different from the background noise. This was done by comparing node activities (perturbation experiment logarithmic fold changes) to the corresponding reference node activity variance (wild type data logarithmic fold change variance) using t-tests. Because we had found that the biological variance was larger than the technical noise, only the number of biological replicates was used to estimate the degrees of freedom. To account for the multiple hypothesis testing, we adjusted the overall false detection rate (FDR) [4]. For the analysis of the *Saccharomyces cerevisiae* genome-wide knockout library, we followed the recommendation by Kemmeren et al. (supplementing information) to exclude several genes from further analysis. Furthermore, we excluded all Pseudogenes and dubious ORFs listed on yeastgenome.org [5]. Because we had found that the distribution of normalized wild type experiment logarithmic fold changes can be well described by a t-distribution with 11 degrees of freedom, this number was used as the degrees of freedom for the t-tests instead of the number of biological wild type experiments. Although the microarray design used by Kemmeren et al. could be used to disentangle technical and biological noise, the biological noise clearly dominated the technical noise. Hence, we did not disentangle the variances for our analyses but rather used the overall variance. Nodes that were significantly affected in at least one perturbation experiment were kept in the network; all other nodes were removed from further analysis.

#### 2.3 Identification of significant links

Inferable links were tested for significance using z-tests. The estimators of the link strength’s variances were found by bootstrapping over node activity data (described below). The false discovery rate (FDR) was adjusted to 5 % to account for multiple hypothesis testing [4].

#### 2.4 Bootstrapping

Residual bootstrapping was used (this was done to get a speed-up compared to parametric bootstrapping, in which residuals are drawn de novo from a distribution rather than used multiple times but in randomized order). First, residuals were calculated from the reference node activity data (wild type data logarithmic fold changes) and then added to the average node activity values of each perturbation experiment. For this addition step, we followed two approaches as explained in the following.

##### 2.4.1 Utilizing correlations among reference node activities for analysis of yeast deletome data

Because we had found that wild type experiment logarithmic fold changes are highly correlated in the yeast deletome data, we sought to maintain the correlations among the reference node activity residuals to improve inference performance.

##### 2.4.2 Minimizing the effect of random correlations

Because we did not simulate correlated measurement noise in our simulated data, the effect of random correlations among the reference node activities was sought to be minimized. This was achieved by shuffling the the order of reference experiments among the nodes.

##### 2.4.3 Residual bootstrapping procedure illustrated

Both bootstrapping approaches are illustrated in the following example, in which it is described how residual bootstrapping is performed on the node activity data for one perturbation experiment. (Residual bootstrapping of reference node activity data is analogous). The node activity data for *N* reference experiments of a two-node network with nodes *i* and *j* is represented as column vectors 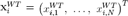 and 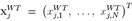, and node activity data for a single perturbation experiment with two replicates is represented as 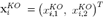 and 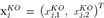. The residuals for the nodes are calculated to be 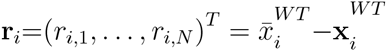 and 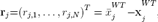, with 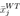 and 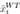 the mean of the corresponding reference node activities. To leave correlations among reference node activities intact, we have the following expression for an arbitrary bootstrap sample for the perturbation node activities: 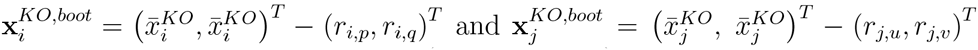, with *p* and *q* being random (possibly equal) integers between 1 and *N*, and 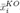 and 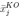 Othe average node activities of the perturbation experiment. To destroy correlations among reference node activity residuals, the residuals are shuffled randomly across bootstrap samples. We have the following expression for an arbitrary bootstrap sample for the perturbation node activities: 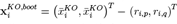 and 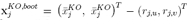 with *p*, *q*, *u* and *v* being random (possibly equal) integers between 1 and *N*.

#### 2.5 Clustering

Nodes were grouped into clusters if they were not sufficiently linearly independent. The following steps were followed:

1. All node activities were normalized to unit reference node activity (wild type data logarithmic fold change) variance.
2. A (*P* × *N*) node activity matrix was formed by merging experimental replicates (here, *P* denotes the number of experiments and *N* denotes the number of genes). Replicates (biological and technical) were merged by averaging and multiplication by the square root of the number of biological replicates.
3. For each pair of nodes *k* and *l*, a test statistic *T*_*kl*_ was compared to a 5% FDR cutoff. The test statistic was calculated as the square of the smaller singular value of the *P* × 2 matrix consisting of those two columns of the node activity matrix which corresponded to the nodes *k* and *l*. For simplicity, we assumed that the reference node activities were Gaussian distributed and that the node interactions were sufficiently linear. Then, under the Null Hypothesis of linear dependence, the test statistic approximately follows a Chi-square distribution with P degrees of freedom (the expected value of the 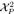 distribution over-estimates the expected value of the small eigenvalue, but this is negligible if sufficiently high confidence is claimed to reject the null hypothesis). An FDR of 5 % was chosen to define a significance cutoff.
4. Clusters were finally formed by grouping unperturbed nodes with perturbed nodes on which they were linearly dependent. If an unperturbed node was linearly dependent on multiple perturbed nodes, this node was grouped with the perturbed node onto which it was the most linearly dependent. Remaining unperturbed nodes were grouped with other unperturbed nodes if they were not sufficiently linearly independent.
5. For each cluster, either the perturbed node, or, if no perturbed node existed in the cluster, the node with the largest signal-to-noise ratio was chosen to be the cluster representative, and the corresponding node activity data was used for further analysis.

Links between clusters were found by inferring links between cluster representatives. Links within clusters can only be inferred within two-node clusters if there is exactly one perturbed node and if the unperturbed node is the only node affected by the perturbed node. Note that all perturbation experiment node activities were used to create the (*P* × *N*) node activity matrix and that no bootstrapping was done.

#### 2.6 Subnetwork method

To infer a link from a node *j* onto a target node i using the subnetwork method, the network is temporarily limited to only those nodes that are significantly affected when node *j* is perturbed (the cutoffs used here were the same ones that were used for identifying the significantly affected nodes). This means that the inference of several links is skipped, and the corresponding link strengths are set to zero and are not included in the calculation of the FDR to determine a cutoff for link significance.

### 3 Applications

### 3.1 Analysis of yeast deletome data

#### 3.1.1 Inference of GAL network

The network was limited to all genes that could in principle be affected by perturbations of the following nodes: GAL3, GAL4, GAL80 and MIG1. A false discovery rate of 10^-3^ was used as a threshold for the identification of significantly affected nodes. With this threshold, 44 nodes remained in the network, that were then clustered. Only clusters that contained at least one of the above mentioned nodes were kept for the graph shown in figure 2d.

#### 3.1.2 Lists of inferred links (consensus stringent.xlsx and consensus less stringent.xlsx)

To generate a list of the most significant links inferred from the whole yeast deletome data, a consensus list was created that reflects the links inferred from the data by using two different methods of bootstrapping (see also the section on bootstrapping). The reason for this is that we are lacking a sophisticated model describing the process that generates biological noise. A consensus list should thus represent a more reliable model for link inference, as only the most significant links that were found by all methods survive the selection process. We created two different consensus lists (consensus stringent.xlsx and consensus_less_stringent.xlsx), each corresponding to a certain cutoff to select significantly affected nodes. That is, for the most stringent cutoff, the network size was smaller than for the less stringent cutoff because less nodes were found to be significantly affected by perturbations. The cutoffs correspond to false discovery rates of 10^-8^ and 10^-6^. The covariance matrix was calculated from all available node activity data for the significantly affected nodes. The following methods for residual bootstrapping were used (see also the section on bootstrapping) to create two separate link lists:

1. Correlations between residuals were left intact.
2. Correlations between residuals were destroyed.

The lists were merged by keeping only links that appeared in both lists and by then keeping the (maximally 500) most significant links with the highest average Z-score. Gene descriptions from the Saccharomyces Genome Database [5] were added to the list to allow easier investigation. We found that, among the most significant links, there are links whose source and target nodes are either adjacent on the genome or which are paralogs. These links are most likely false positives that reflect inspecificities of the gene deletion process used to generate the yeast deletion collection [2].

#### 3.1.3 List of inferred hub nodes (consensus_stringent_hubs.xlsx)

From the consensus link list with the stringent cutoff for the selection of significantly affected nodes, we selected the 10 nodes with the highest number of significant outgoing links (significance cutoff: 5 % false discovery rate). The mean and median link strengths of the outgoing links were calculated for each of these 10 hub nodes. Gene descriptions from the Saccharomyces Genome Database [5] were added.

### 3.2 Simulations

#### 3.2.1 Effect of noise-induced bias on relative link strength (suppl. figure 1d)

To show that network inference methods are biased towards measurement noise, we simulated node activity data for the limiting case of infinitely many replicates. In that case, all parameters are estimated perfectly because uncertainty resulting from a limited number of observations are averaged out. The covariance matrix of node activities was simulated according to the following formula:

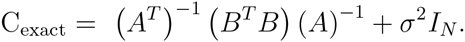

Where *B* is a diagonal matrix with element *b*_ii_ = 1 if node *i* is perturbed. The signal-to-noise ratio was adjusted by changing the value of *σ*^2^.

#### 3.2.2 Comparison of simple, clustering and subnetwork methods (figure 2c)

The Inferability curves represent averages over 4 different networks and 12 perturbation samples. The network size was 60 nodes. Gene expression data was simulated from scale free networks and a non-linear model (as described below).

#### 3.2.3 Receiver Operating Characteristic (ROC) curves (figure 2b)

##### Simulation setup

The ROC curves were averaged (as described below) over 24 perturbation samples and 4 network structures. The network size was 300 nodes, and 25 % of nodes were randomly perturbed in each perturbation sample. The noise-to-signal ratio was adjusted to 10 %. Data was generated with both GeneNetWeaver [3] and scale-free networks with a non-linear model to generate stationary data. For each perturbation sample, only data corresponding to significantly perturbed nodes was used to calculate the covariance matrices (1% FDR), the only exception being Lasso regression applied to the data of all nodes. All methods (except Lasso applied to the data of all nodes) received almost the same covariance matrix C as input: because of the randomness of bootstrap sampling, small numerical differences between the covariance matrices that PRC and the other methods received may have occurred. Binary classification was done solely based on links pointing away from perturbed nodes because only those links are potentially inferable. This is similar to the method described by Siegenthaler and Gunawan [6]. To infer a link onto a certain node, the data corresponding to a perturbation of that node was removed prior to calculating the covariance matrix. Because both Lasso regression and the subnetwork method set certain link strengths to exactly zero, it would be impossible to smoothly reach a false positive rate of 1. This is why we set all link test statistics that were set to zero to random values smaller than the smallest non-zero link test statistic. When creating the ROC curve, this procedure corresponds to random guessing for all links that were set to exactly zero.

##### ROC curve averaging

In the following, the false positive rate is denoted by FPR and the true positive rate is denoted by TPR. To generate an averaged ROC curve, the tuples (FPR, TPR) of the individual ROC curves were assigned to 50 bins according to the FPR of each tuple. Bins were filled up in a way such that approximately equally many tuples were assigned to all bins. Then, for each bin, the FPR and TPR was calculated. These 50 averaged tuples (FPR, TPR) were used for the averaged ROC curve.

##### ROC curves for Subnetwork method and Lasso

We generated 10 different ROC curves for the subnetwork method, each curve corresponding to a different cutoff for the selection of significantly affected nodes (based on which the subnetworks were created). For Lasso, we generated ROC curves for 10 different regularization coefficients. To model a smooth transition between the cutoffs or regularization coefficients, we interpolated (cubic spline) between the ROC curves in a way such that the resulting hybrid curve had a possibly larger area under the curve than the individual ROC curves. The 10 different cutoffs for selection of significant nodes correspond to the following FDRs: 1.00, 0.89, 0.78, 0.67, 0.56, 0.45, 0.34, 0.23, 0.12, and 0.01. The 10 different regularization coefficients were: 0, 0.001, 0.0018, 0.0032, 0.0056, 0.010, 0.0178, 0.0316, 0.0562, and 0.1000.

##### Comparison of non-regularized inference methods

Test-statistics *T*_*ji*_ corresponding to a link from a node *j* to a node *i* were calculated according to the following formulas. The standard deviations of the test statistics were estimated via residual bootstrap.

1. Regression: 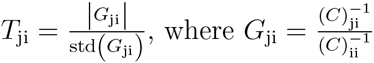.
2. Conditional Mutual Information: 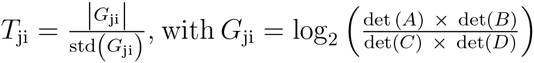. Here, *A, B* and *D* are covariance matrices with rows and columns of certain nodes removed. *A:* node *j* removed. *B:* node *i* removed. *D*: nodes *i* and *j* removed.
3. PRC: 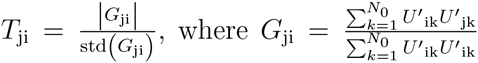, where *N*_0_ is the number of unperturbed nodes and *U*_*i*_ is the i^th^ eigenvector of the covariance matrix. Note that links were not identified as artifacts.

##### Comparison of regularized inference methods

For a fair comparison, we used 10 different regularization coefficients to generate ROC curves for Lasso regression, keeping only the ROC curve corresponding to the regularization coefficient that gave the best performance. The following is a brief derivation of the formulas used for regularized regression using the L_1_ and L_2_ norms. The simulation setup is described afterwards. Consider the following equation, which is essentially equation 1 of the main paper except that perturbations are not restricted to single nodes, which is expressed through the perturbation strength matrix *U:*

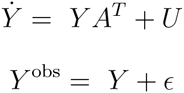

For simplicity, we assumed here that the reference state is 0. Here, Yare the true node activities that are masked by measurement noise e to yield observed node activities *Y*^*obs*^, and the matrix A^T^denotes the link strengths. In the steady state and under the assumption that the measurement uncertainties *ϵ* are Gaussian distributed, a maximum a posteriori (MAP) function of the parameters *A*^*T*^ and *U* given the observed data *Y*^*obs*^ can be defined. The logarithm of this function is:

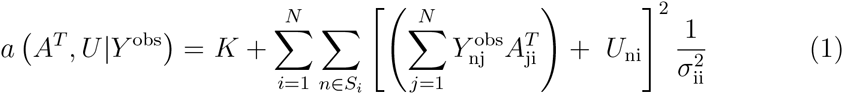

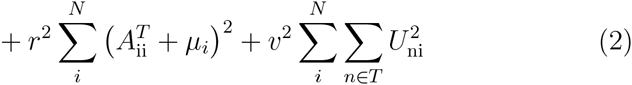

Here, *K* is a normalization constant from the distributions that vanishes upon differentiation. The experiment indices *n* run over the set of all experiments except the ones in which node *i* is perturbed, *S*_*i*_. *N* is the number of nodes; *P* is the number of experiments; 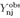 is the matrix of node activities (each row corresponds to a sample and each column corresponds to a node); 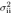 is a variance coefficient and *r*^2^,*v*^2^ are regularization coefficients that correspond to weights of the prior distributions. Note that the first prior distribution is only defined over the diagonal elements of the network matrix *A*. It can be shown that the following estimator maximizes the MAP function:

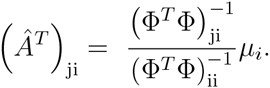

Here, Φ denotes the observed node activities with samples corresponding to a perturbation of node *i* removed (the number of rows of Φ is less than or equal to the number of rows of *Y*^*obs*^, depending on whether node *i* is perturbed). This means that the data may need to be re-organized to infer links onto each node. To arrive at this result, one needs to maximize *a*(*A*^*T*^,*U*|*Y*^*obs*^) with respect to *A*. This is possible because the diagonal elements of the network matrix *A* are regularized, which does not allow for arbitrary combinations of either *A* or *U*. The following steps lead to the above stated solution:

1. Set the derivative of the log-MAP function w.r.t. *U*_*ni*_ to zero: 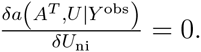
2. Solve for *U*_*ni*_ to obtain the MAP estimator *U*̂_ni_. Plug this expression into formula for log-MAP function.
3. Set this new expression of the log-MAP function w.r.t. 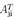 zero: 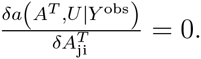
4. Solve for 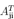 to obtain the MAP estimator 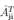. To achieve this, one has to apply the Sherman-Morrison-Woodbury formula and let the regularization coefficient *r*^2^ go to infinity, which essentially sets all 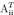 equal *µ*_*i*_.

Note that, for *µ*_*i*_ = *–*1, the set of parameters 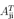 (except for 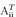, which become equal to *µ*_*i*_) are simply the ordinary least squares (OLS) estimators that minimize the following expression:

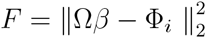

Here, Ω is a matrix that is equal to Φ except that it is missing column *i*, and *β* is a vector that is equal to 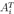 except that it is missing the element equal to 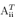. The L_1_ Norm (“Lasso” regularization) and L_2_ Norm (“Tikhonov-Miller” regularization) impose further constraints on this extreme value problem. The corresponding formulas are:

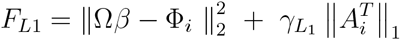

and

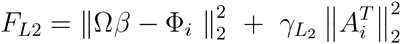

for the L_1_ and L_2_ Norm, respectively.

### 4 Software

For all analyses and simulations, we used MATLAB and Statistics Toolbox and Parallel Computing Toolbox Release R2014b, The MathWorks, Inc., Natick, Massachusetts, United States. Our MATLAB code is available on demand.

